# Structure and functional analyses of vaccinia virus J5 protein reveal distinct determinants for entry-fusion complex assembly and activation

**DOI:** 10.64898/2026.03.08.710424

**Authors:** Hsiao-Jung Chiu, Kathleen Joyce Carillo, Louise Tzung-Harn Hsieh, Yuan-Chao Lou, Chang Sheng-Huei Lin, Der-Lii Tzou, Wen Chang

**Author notes:** Corresponding author: correspondence should be addressed to: Wen Chang, Der-Lii Tzou, Chang Sheng-Huei Lin. These authors contributed equally to this article (Hsiao-Jung Chiu, Kathleen Joyce Carillo, Louise Tzung-Harn Hsieh).

## Abstract

Vaccinia virus enters host cells through a multi-component entry fusion complex (EFC) that is structurally distinct from canonical viral fusion systems. Understanding how vaccinia virus EFC mediates membrane fusion is crucial for elucidating poxvirus entry and identifying potential antiviral targets. Here, we report the solution NMR structure of a truncated ectodomain of vaccinia J5 protein, residues 2-68. Using recombinant vaccinia viruses expressing J5 mutants, we analyzed substitutions in conserved and surface-exposed residues, as well as chimeric constructs between vaccinia J5 and its entomopoxvirus ortholog AMV232. Functional analyses revealed that the conserved P^38^YYCWY^43^ motif is dispensable for EFC assembly but required for membrane fusion activity whereas the flexible region spanning residues 90-110 mediates interactions required for stable incorporation of J5 into the EFC.

**Importance:** Vaccinia virus enters host cells through membrane fusion mediated by a unique multi-component entry fusion complex (EFC) that is distinct from classical viral fusion proteins. Although J5 has been identified as a central component of the pre-fusion EFC, the structural regions of J5 required for membrane fusion remain unclear. Here, we determined the solution NMR structure of the J5 ectodomain and identified two regions, the conserved P^38^YYCWY^43^ motif and residues 90-110, as key determinants of EFC function during vaccinia virus entry.

## Introduction

Vaccinia virus (VACV) is a prototypic member of the family *Poxviridae* and contains a linear, double-stranded DNA genome of approximately 194.7 kilobase pairs, encoding more than 200 open reading frames (1, 2). VACV has served as a model virus for studying poxvirus molecular biology and virus-host interactions. Historically, vaccinia virus was used as the live vaccine that led to the global eradication of smallpox caused by variola virus (3). More recently, it has also been employed as a vaccine platform to prevent infection by emerging orthopoxviruses such as monkeypox virus (Mpox) during recent outbreaks (4).

VACV has a broad host range and completes the virus life cycle in the cytoplasm of host cells (5). Upon infection and subsequent core uncoating, viral factories form in perinuclear regions of the cytoplasm and further assemble into mature virus (MV) containing a single membrane derived from the endoplasmic reticulum (6–9). A portion of these infectious particles is then transported to the Golgi cisternae, where additional membrane wrapping occurs (10, 11), and the virus subsequently egresses as a double-membrane extracellular virus (EV) (12–14). Although MVs and EVs mediate distinct modes of transmission and enter host cells via different mechanisms (15–17), both forms of viral particles utilize an identical multiprotein (18), highly-conserved entry-fusion complex (EFC) for membrane fusion between viral and host membranes. The EFC contains 11 component proteins: A16 (19), A21 (20), A28(21), F9 (22), G3 (23), G9 (24), H2 (25), J5 (26), L1 (27), L5 (28) and O3 (29). Unlike the vast majority of viruses that utilize a single viral protein to engage cellular receptors and subsequently trigger membrane fusion through conformational changes (30); poxviruses instead employ multiple proteins. At least four viral proteins mediate attachment to glycosaminoglycans or receptors on the cell surface (31–34). In addition, each EFC component is pivotal for the viral infectivity, EFC assembly, and membrane fusion activity, although individual proteins may function at distinct stages of the fusion process (35).

Structures of individual EFC components have been determined, including A21 (36), A28 (37), F9 (38), H2 (39), L1 (40), the subcomplexes A16/G9 (41), and G3/L5 (42), and most recently the whole EFC (43). However, none of these structures or protein sequences exhibit detectable homology to well-characterized fusion proteins (44) of the class I, II or III (45, 46), suggesting that VACV employs a distinct membrane fusion mechanism.

Among the VACV EFC components, two structurally conserved families have been identified: A16/G9/J5 and F9/L1 (47, 48). These families share conserved cysteine positions and disulfide bonding patterns, which are widely distributed across the *Poxviridae* family (48), suggesting a conserved fusion mechanism during evolution. J5 is the shortest protein within the evolutionarily conserved A16/G9/J5 family (26). Attempts to generate a J5 knockout virus were unsuccessful, indicating that J5 is essential for virus growth in cells (49). J5 associates with G9 and A16 via the conserved PXXCW and Delta motifs shared among the three orthologs, thereby stabilizing the central core in the cryo-EM structure of the EFC (43). However, the specific residues of J5 that regulate membrane fusion remain poorly defined. In this study, we determined the solution NMR structure of a truncated J5 ectodomain protein. Through mutagenesis and recombinant virus construction, we identified critical residues in J5 required for membrane fusion and EFC assembly, providing new insights into the functional architecture of the vaccinia virus entry-fusion complex.

## Materials and Methods

### VACV J5 plasmid construction, protein expression and purification

The cDNA encoding the soluble fragment of the vaccinia virus J5 protein (residues 2-68; tJ5) was cloned into the pETDuet-1 vector with an N-terminal His₆-SUMO (Smt3) fusion tag under the control of the T7 promoter. The construct was transformed into *E. coli* BL21(DE3) cells for protein expression. For isotopic labeling, uniformly labeled ^15^N/^13^C protein was expressed in M9 minimal medium containing 1 g/L ^15^NH_4_Cl (Sigma-Aldrich) as the sole nitrogen source and 1 g/L uniformly labeled ^13^C_6_ glucose (Cambridge Isotope Laboratories) as the sole carbon source. The medium was supplemented with 2 mM MgSO_4_, 0.1 mM CaCl_2_, 1 mg/L thiamin, and 1 mg/L biotin. Cells were initially grown overnight in 30 mL of LB medium and then transferred to 1 L of M9 medium after centrifugation to remove residual LB medium. Protein expression was induced by adding 0.5 mM isopropyl-β-d-thiogalactoside (IPTG) when the culture reached an optical density at 600 nm (OD_600_) of 0.6. Cells were incubated for 18 h at 16 °C and harvested by centrifugation at 3,000 × g for 30 min at 4 °C.

Cell pellets were resuspended in lysis buffer containing 20 mM Tris (pH 8.0), 50 mM NaCl, 1 mM phenylmethylsulfonyl fluoride (PMSF), 1 mM 1,4-dithiothreitol (DTT), and lysozyme until homogeneous. Cell lysis was performed by sonication on ice. The lysate was cleared by centrifugation at 30,000 × g for 1 h at 4 °C and filtered through a 0.45 μm filter. The clarified supernatant was incubated overnight at 4 °C with 1 mL Ni^2+^-NTA affinity resin (Bio-Rad) packed into an empty PD10 column. The resin was washed three times with 10 column volumes of 20 mM Tris (pH 8.0) containing 500 mM NaCl and 20 mM imidazole. The protein was then eluted with 20 mM Tris (pH 8.0), 500 mM NaCl, 500 mM imidazole, and 1 mM DTT. To remove the His_6_-SUMO tag, Ulp1 protease (50) was added during overnight dialysis in 20 mM Tris (pH 8.0) and 50 mM NaCl at 4 °C. Untagged tJ5 protein was subsequently separated using Ni-NTA resin (Bio-Rad). The protein sample was concentrated to 0.5 mM using an Amicon Ultra Centrifugal Filter with a 3-kDa molecular weight cutoff (Millipore). The concentrated protein was then exchanged into NMR buffer containing 10 mM phosphate (pH 6.5) and 137 mM NaCl and 2.7 mM KCl. Protein concentration was determined by absorbance at 280 nm (ε280 = 13,325 M^-1^ cm^-1^), and purity was confirmed by 15% SDS-polyacrylamide gel electrophoresis (SDS-PAGE).

### Nuclear magnetic resonance (NMR) spectroscopy and structural determination

All NMR spectra were recorded at 298 K and pH 6.5 using a Bruker AVANCE 600- or 800-MHz spectrometer equipped with a 5-mm triple-resonance TXI cryogenic probe with a shielded Z-gradient. Samples of 0.5 mM ^15^N/^13^C-labeled tJ5 in 1X PBS buffer (90% H_2_O and 10% D_2_O at pH 6.5) were loaded into 5-mm Shigemi NMR tubes for the experiments.

Sequential backbone resonance assignments were obtained through independent connectivity analysis of NHCACB, CBCA(CO)NH, HNCO, and HN(CA)CO experiments. NOE restraints were derived from 3D ^13^C- and ^15^N-edited NOESY-HSQC spectra with a mixing time of 150 ms. Backbone assignments and NOE resonances were analyzed using Topspin 4.3.0 (Bruker) and CARA 1.8.4 (51). NOE cross-peaks were categorized as very weak, weak, medium, or strong based on signal intensity. Hydrogen bond restraints were identified by monitoring slow HN exchange with solvent D_2_O. Dihedral angle restraints (ϕ and Ψ) were predicted from chemical shift data using the TALOS+ web server (52). For the final set of protein structure calculations, 100 structures of tJ5 were generated using the XPLOR-NIH 3.4 (53, 54) with a standard simulated annealing protocol, including high-temperature dynamics at 1000 K followed by cooling to 100 K. The 20 lowest-energy structures were selected for water refinement using the AMPS-NMR web portal (55) and a standard restrained molecular dynamics protocol implemented within the AMBER99SB-ILDN force field (56). Refinement employed a generalized Born solvent model with an ionic strength of 137 mM NaCl, and a 10 Å TIP3P water box. The final ensemble of 10 structures contained no distance or dihedral angle violations exceeding 0.5 Å or 5°, respectively. Structural quality was assessed using the Protein Structure Validation Suite (PSVS) (57). Protein structure figures were generated using the Chimera 1.18 (58). Statistics for the final structure are summarized in Table 1. The coordinates for the 10 conformers of tJ5 have been deposited in the Protein Data Bank under accession code 8WT5.

**Table 1.**
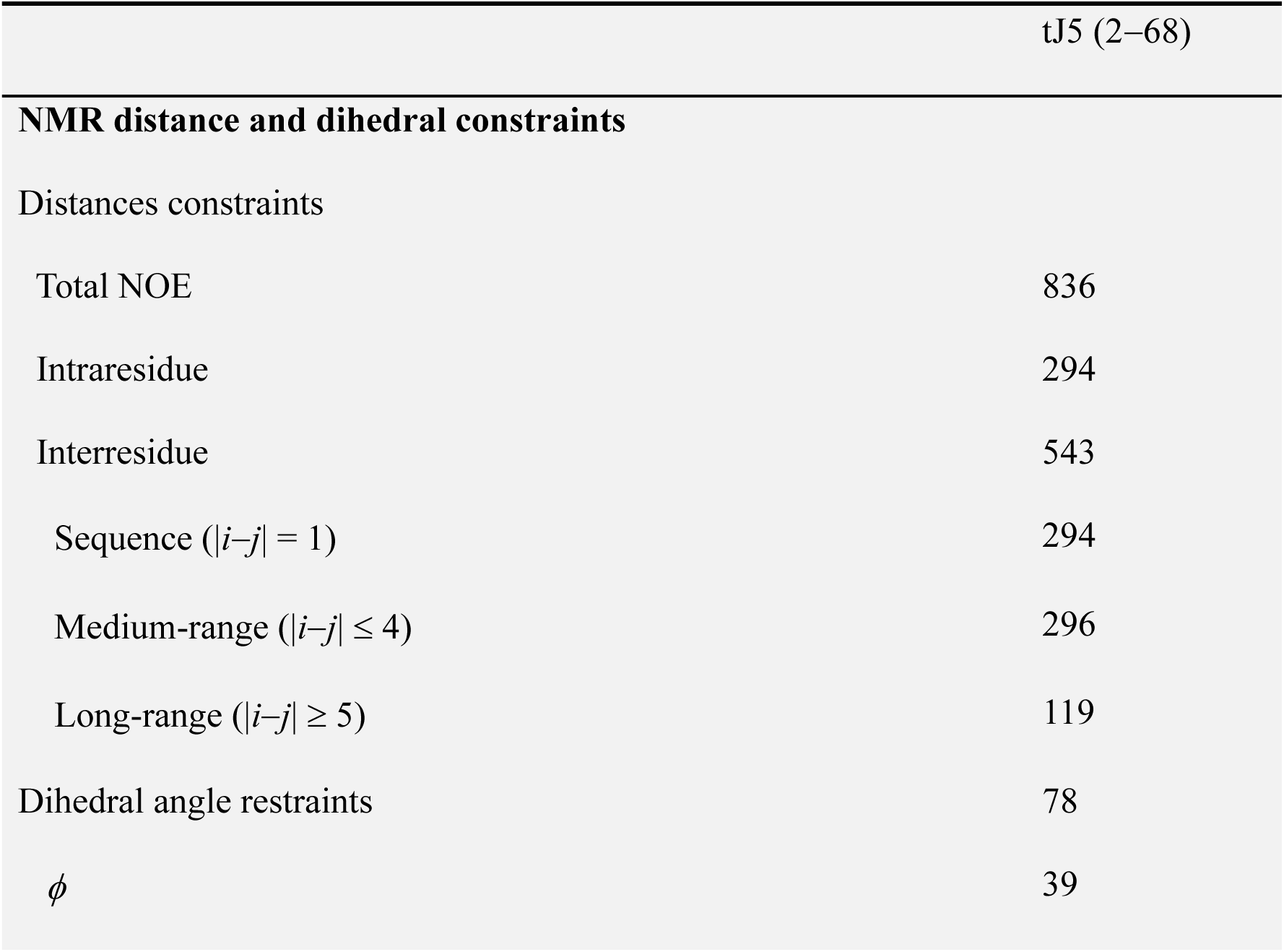
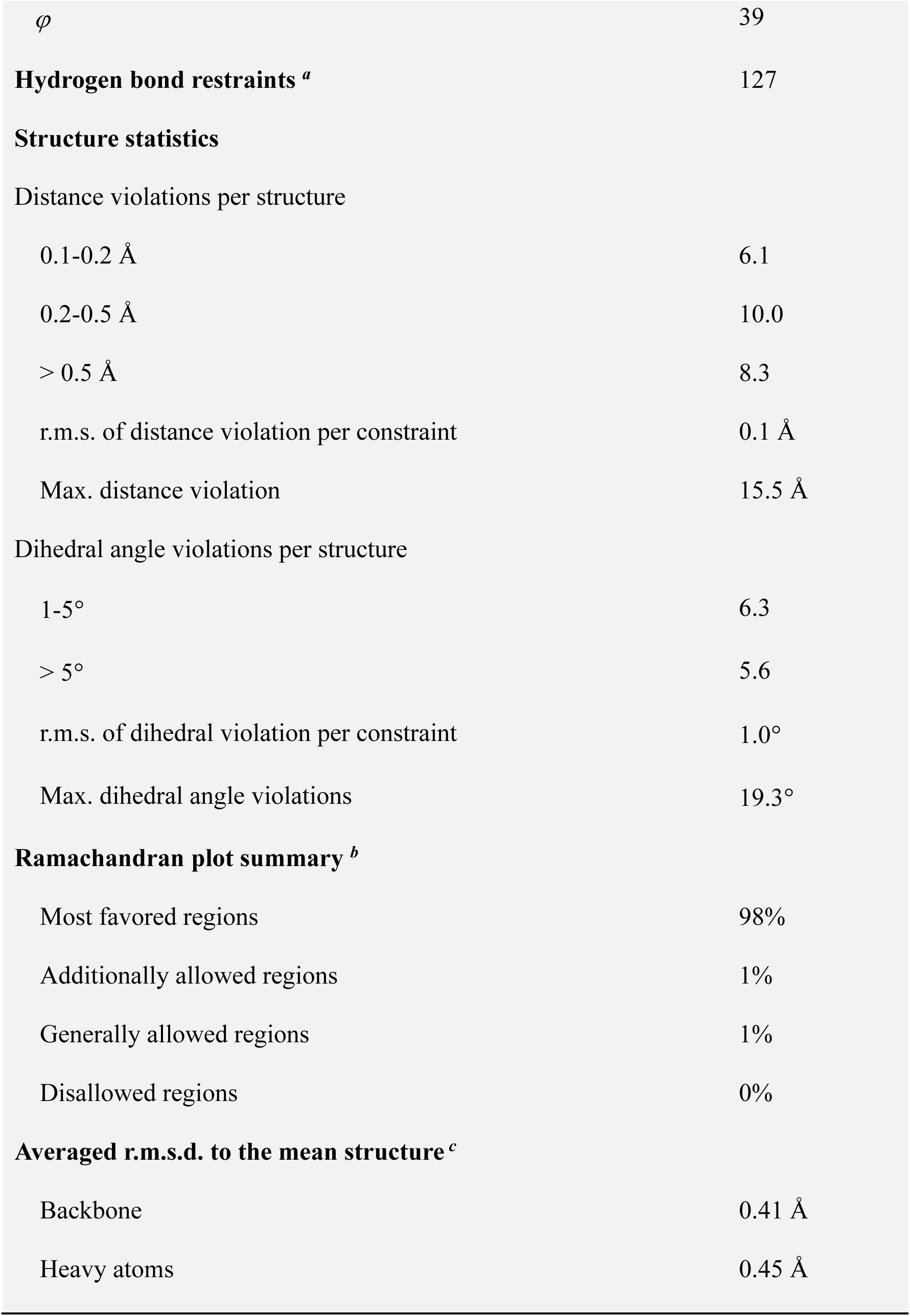
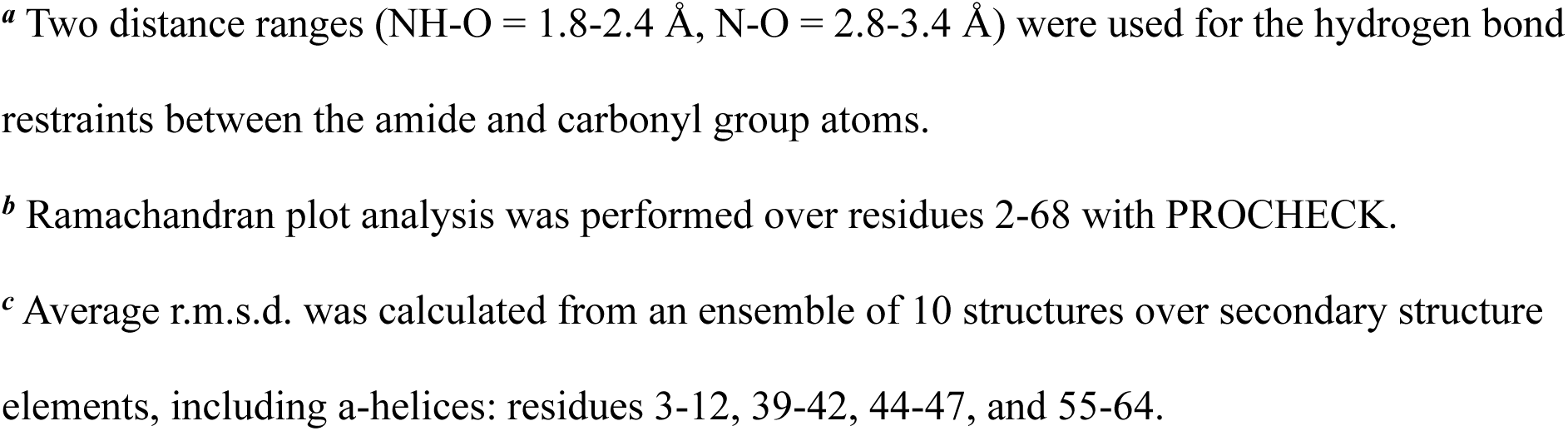
NMR and refinement statistics for vaccinia virus fusion protein tJ5.

### Cells, viruses and reagents

BSC40, CV-1, GFP- and RFP-expressing HeLa cells were cultured in Dulbecco’s modified Eagle’s medium (DMEM) supplemented with 10% fetal bovine serum (HyClone), 100 units/mL Penicillin and 100 µg/mL Streptomycin (Gibco) as previously described (59). The Western Reserve (WR) strain of vaccinia virus and recombinant viruses constructed in this study were propagated in BSC40 cells. Restriction enzymes were purchased from New England Biolabs. Fixative and staining solutions for electron microscopy analysis were obtained from Electron Microscopy Sciences.

### Bioinformatic analyses of poxviral J5 orthologues

Orthologs of the vaccinia J5 protein from representative members of the *Poxviridae* family were retrieved from GenBank with the following gene accession numbers: YP_232979.1 (VACV), NP_570485.1 (camelpox virus), NP_619893.1 (cowpox virus), NP_671599.1 (ectromelia virus), NP_536516.1 (monkeypox virus), YP_009282790.1 (skunkpox virus), YP_717406.1 (taterapox virus), NP_042126.1 (variola virus), YP_009281844.1 (volepox virus), NP_039099.1 (fowlpox virus), YP_009177120.1 (turkeypox virus), YP_001293261.1 (goatpox virus), YP_004821430.1 (yokapox virus), QGT49348.1 (crocodilepox virus), YP_008658487.1 (squirrelpox virus), NP_051781.1 (myxoma virus), NP_044029.1 (molluscum contagiosum virus), YP_009480606.1 (sea otterpox virus), NP_957964.1 (bovine papular stomatitis virus), YP_009112794.1 (parapoxvirus), YP_003457360.1 (pseudocowpox virus), NP_957832.1 (orf virus), YP_009268788.1 (pteropox virus), AKR04183.1 (sapmonkeypox virus), NP_938325.1 (yabapox virus), YP_009001515.1 (anomala virus), YP_009408025.1 (eptesipox virus), and NP_065014.1 (Amsacta moorei entomopoxvirus, AMV). Multiple sequence alignments were generated using MAFFT v7.453 with parameters --maxiterate 1000 --globalpair, and visualized with Jalview v1.8.3. Residue conservation was color-coded as follows: yellow (identical), blue (>0.5 conservation), and green (>0.2 conservation). Protein sequences of VACV J5 and AMV232 were further aligned using the ClustalW algorithm in MacVector (v16.0.10) with identical residues shaded in gray.

### Protein structure prediction with AlphaFold2

The swap mutation protein structures were predicted using AlphaFold2 (https://reurl.cc/96anZj). Sequences searches were performed using MMseqs2 (mmseqs2_uniref_env) in unpaired/paired mode. For each protein, the predicted model with the highest overall confidence (pLDDT >70) was selected for analysis. The mean pLDDT score for each model is shown in the corresponding figure.

### Structural alignment and residue interaction analysis

Protein structural similarity was analyzed in UCSF Chimera (version 1.18). Structural superimposition was performed using Matchmaker tool, which first generates a sequence alignment followed by structural fitting of aligned residue pairs. Default alignment parameters were applied, including the Needleman-Wunsch alignment algorism, with alignment score calculated using a weighting of 70% residue similarity based on the BLOSUM62 substitution matrix and 30% secondary structure similarity. Structural fitting was performed using Cα atoms of aligned residue pairs. Root-mean-square deviation (RMSD) values were calculated using the Match-Align function following structural superposition. Residue interaction analysis of EFC (PDB 9UZO) was performed using the Residue Interaction Network Generator (RING; https://ring.biocomputingup.it/) and PyMOL to identify intermolecular contacts.

### Construction of J5 mutant plasmids and recombinant vaccinia viruses

VACV WT and mutant *J5L* ORFs were synthesized (GenScript Inc.) with *Bst*Z17I and *Mfe*I sites at their 5’ and 3’ ends and encode the chimeric proteins listed in the table below.

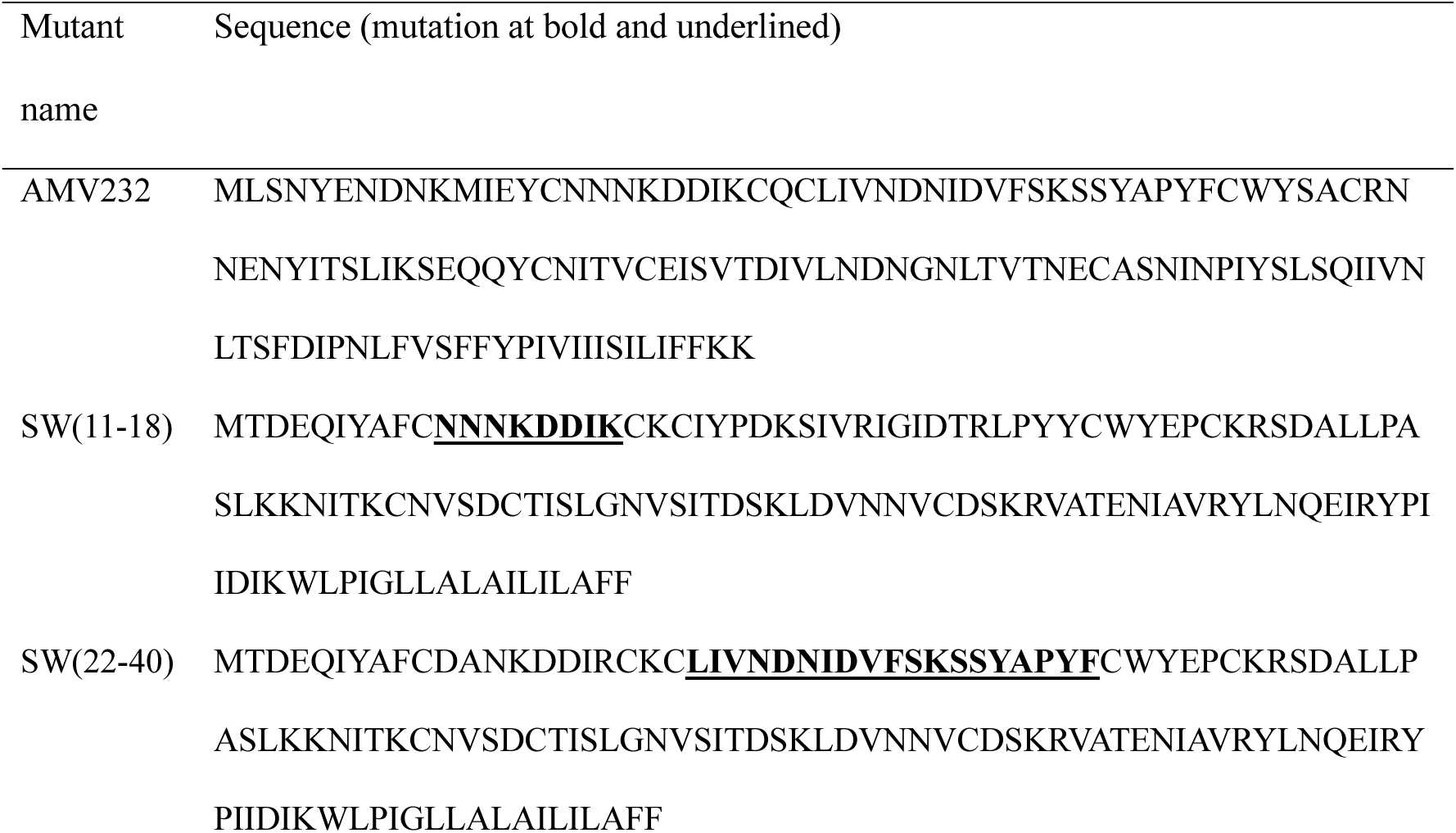
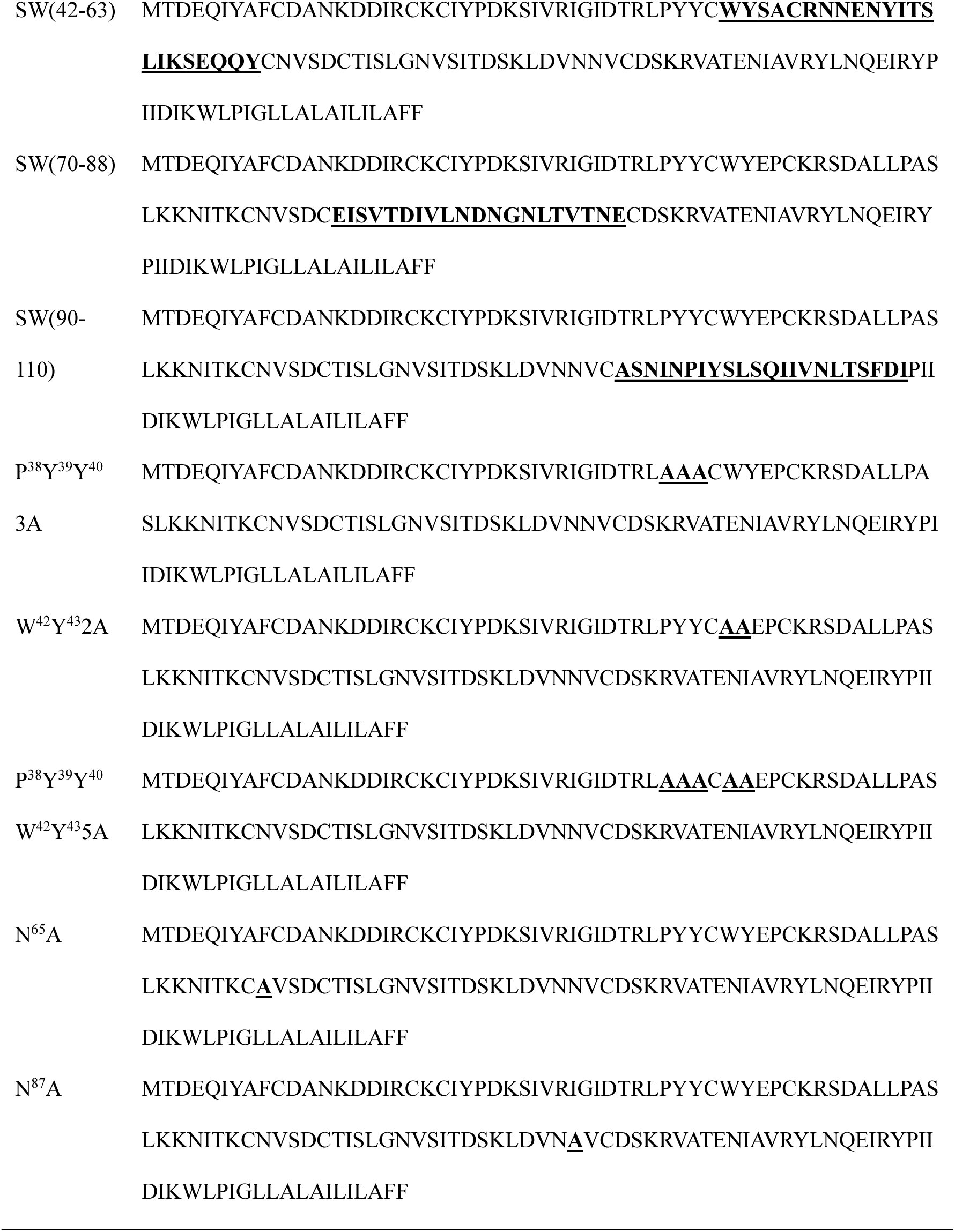

The Eco*Gpt* cassette driven by the p7.5k promoter was constructed into a plasmid containing the 5’ and 3’ flanking sequences of the *J5L* locus in the viral genome as described below. A 1-kb flanking fragment containing partial J6R and J5L promoter sequence was amplified by PCR using WR genomic DNA as template and cloned into pBlueScript KS(-) using primer 5’-TAGGGCGAATTG<u>GGTACC</u>AACGGTGATAGATGTACT-3’ and 5’- GAATTCGATATC<u>AAGCTT</u>GTATACCTTCGGTTCTTT-3’ (the *Kpn*I and *Hind*III sites were underlined). Then, a mutation was introduced to disrupt a *Bst*Z17I site using primers 5’-AATTCATTGGTTTCTTTGGGAATACTATCTATCCAAAAACT and 5’-AGTTTTTGGATAGATAGTATTCCCAAAGAAACCAATGAATT-3’. The Eco*Gpt* cassette and 1-kb flanking fragment containing *J4R* ORF and partial *J3R* were generated by PCR using primers 5’-CAAGCTTGATATCGAATTCAATTGGATCACTAATTCCAAACC-3’ and 5’-TATTTCAGTCGCTGATTAATCTAGCGACCGGAGATTGGCGG-3’, 5’- CCGCCAATCTCCGGTCGCTAGATTAATCAGCGACTGAAATA-3’ and 5’-TGGAGCTCCACCGCGGTGGCGGCCGCCCGTTTCCAGATCAATGGAT-3’, respectively, followed by assembly with linearized pBlueScript-J6 using NEBuilder. Then, *J5L* ORF was amplified by PCR using primers 5’-TAT<u>GTATAC</u>AATCAAATTTCCCTTTTTA-3’ and 5’- TAT<u>CAATTG</u>GTAGAGATGAGATAAGAA-3’ or obtained by double digestion with *Bst*Z17I and *Mfe*I and subsequently subcloned into pBlueScript J6R-gpt-J4R-J3. In several cases, *J5L* mutants were generated from the *J5L* WT plasmid using the QuickChange Lightning site-directed mutagenesis kit (Agilent Technologies) with the primers listed in the table below.

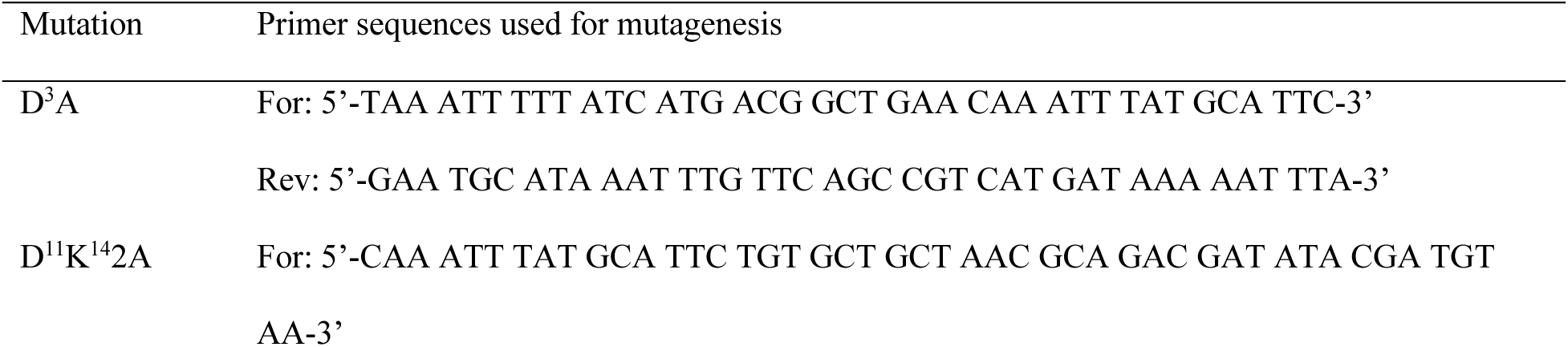
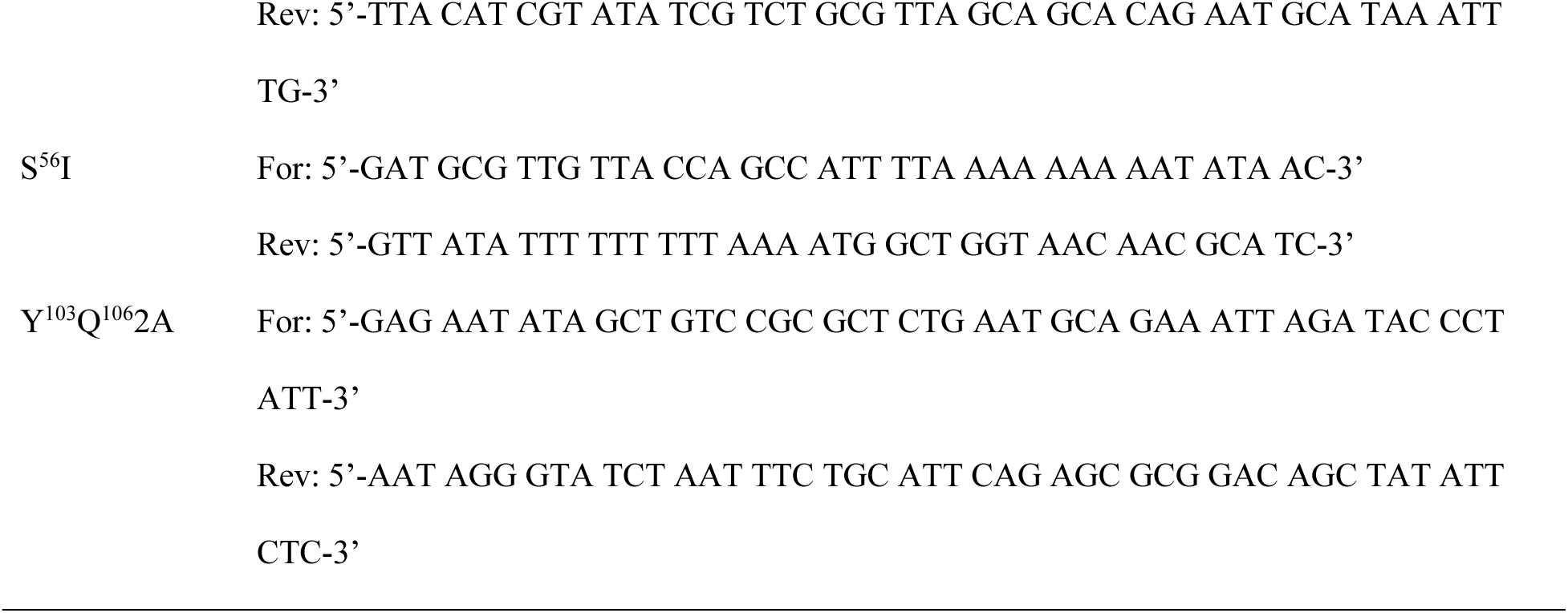

All the constructed *J5L* WT and mutant plasmids were linearized and individually transfected into WR-GS-infected CV-1 cells and harvested at 24 h post-transfection. Recombinant viruses were subsequently isolated by at least three rounds of plaque purification in 1% agar under *gpt* selection (25 µg/mL mycophenolic acid, 250 µg/mL xanthine, and 15 µg/mL hypoxanthine) as described in the established protocol (60).

### Determination of J5 mutant virus infectivity

BSC40 cells were seeded in six-well plates and infected with J5 WT or J5 mutant recombinant viruses at a multiplicity of infection (MOI) of 5 PFU per cell. After 1 h of adsorption at 37 °C, cells were washed with PBS and cultured in fresh growth medium for 24 h before harvesting for plaque assays and immunoblot analysis. All experiments were performed independently three times.

### MV-triggered cell-cell fusion at acid pH (Fusion-from-without)

VACV MV-mediated cell-cell fusion assays were performed as previously described (37, 39, 59). In brief, HeLa cells expressing GFP or RFP were mixed at a 1:1 ratio and seeded at 3×10^4^ cells/well in a 96-well plate. The following day, cells were pretreated with 40 μg/mL cordycepin (Sigma) for an hour and subsequently incubated with J5 WT recombinant virus at a MOI of 100 PFU/cell or an equal amount of lysates of each J5 mutant recombinant virus in 50 μl at 37 °C for 60 min. Cells were subsequently washed, treated with neutral (pH 7.4) or acidic (pH 5.0) PBS at 37 °C for 3 minutes, washed with growth medium, and replaced with fresh medium containing cordycepin as described above. Cell fusion was monitored every 30 minutes using ImageXpress Confocal HT.ai High Content Imaging system (Molecular Devices) until 3 hours postinfection. Fusion efficiency was quantified using Fiji software as follows: (surface area of GFP^+^RFP^+^ double-fluorescent cells/surface area of single-fluorescent cells) × 100%, as previously described (37, 39, 59). All experiments were performed independently three times.

### Immunoblot analysis

Protein samples were denatured in SDS-containing sample buffer at 95 °C for 10 min and analyzed by immunoblotting. Samples were separated by SDS-PAGE and transferred onto 0.45 µm nitrocellulose membranes (Bio-Rad). Membranes were probed with polyclonal antibodies against A16, A21, A28, D8, G3, G9, H2, J5, and L5 as previously described (37).

### Coimmunoprecipitation

Confluent BSC40 cells were infected with recombinant J5 WT or mutant viruses at an equivalent MOI of 2 PFU per cell and incubated at 37 °C for 24 h. Cells were washed with PBS, harvested, and lysed in ice-cold buffer containing 20 mM Tris (pH 8.0), 200 mM NaCl, 1 mM EDTA, and 1% NP-40 supplemented with protease inhibitors (1 mM phenylmethylsulfonyl fluoride, 2 µg/mL aprotinin, 1 µg/mL leupeptin, and 0.7 µg/mL pepstatin). After centrifugation to remove cell debris, 250 µg of clarified lysate was incubated with anti-J5, anti-G9, or anti-A28 antisera and protein A-conjugated beads (GE Healthcare). The samples were rotated at 4°C for 16 h, centrifuged, washed five times with lysis buffer, and analyzed by SDS-PAGE and immunoblotting as previously described (37).

### Transmission electron microscopy (TEM) analysis

The TEM analyses were performed as previously described (61). In brief, confluent BSC40 cells were seeded onto poly-L-lysine-coated plastic slides and individually infected with J5 WT or mutant viruses at a MOI of 2 PFU per cell for 24 h. Cells were fixed in 2.5% glutaraldehyde, postfixed with 1% osmium tetroxide, and contrasted with 1% uranyl acetate. After serial ethanol dehydration, samples were embedded in Epon resin and sectioned for transmission electron microscopy (TEM). Samples were imaged on a Talos L120C transmission electron microscope equipped with a CETA 16 4k x 4k CMOS camera (Thermo Scientific).

### Statistical analysis

Statistical analysis was performed using GraphPad Prism v10 (GraphPad Software, San Diego, CA). Data are presented as mean ± standard deviation (SD). Statistical significance was assessed using a two-tailed Student’s *t* test. For comparisons with WT controls, the *P* value was adjusted using “p.adjust” function with the false discovery rate (FDR) method in R v4.5.1. Adjusted *P* values of < 0.05 were considered statistically significant. Significance levels are indicated as follows: *, *P* < 0.05, **, *P* < 0.01; ***, *P* < 0.001 and ****, *P* < 0.0001.

## Results

### Nuclear magnetic resonance (NMR) structure of truncated vaccinia virus fusion component J5 (tJ5)

We generated a bacterial expression plasmid encoding the ectodomain of vaccinia J5 protein (residues 2-68), designated tJ5, fused to a hexa-histidine tag at the N-terminus (Fig. 1A). The tJ5 protein was purified using Ni²⁺ affinity chromatography (Fig. 1B). Using ^15^N and ^13^C isotope labeling, we determined the backbone resonance assignments of tJ5 by three-dimensional heteronuclear NMR spectroscopy, as shown in the ^1^H-^15^N HSQC spectrum (Fig. S1). The 2D HSQC spectrum of isotopically labelled tJ5 displayed well-resolved resonances for most non-proline residues, suggesting that the protein is well-folded in solution. Secondary structure was predicted based on ^13^C_α_ chemical shift propensity analysis, which revealed six α-helices connected by loop regions within the N-terminal ectodomain (residues 2-68).

**Figure 1.**
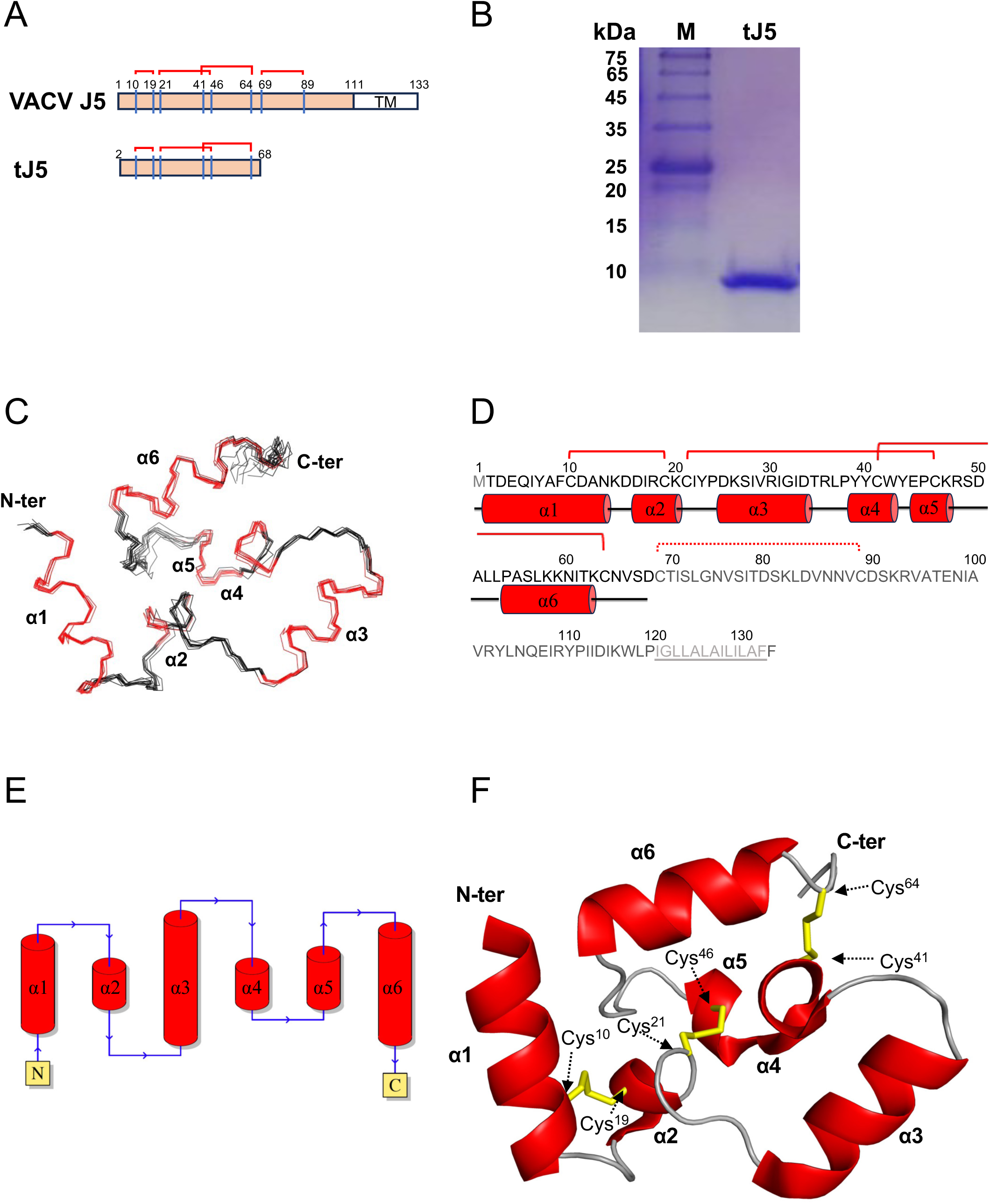
NMR structure determination of the vaccinia virus J5 ectodomain (tJ5, residues 2-68). (A) Schematic representation of the full-length J5 protein (residues 1-133) showing the transmembrane domain (TM; residues 111-133) and the truncated soluble construct (tJ5; residues 2-68) used for structural determination. (B) SDS-PAGE analysis of purified recombinant tJ5 protein stained with Coomassie blue. M, protein markers (kDa). (C) An ensemble of the 10 lowest-energy NMR structures of tJ5. The ribbon is colored with α-helices in red and unstructured regions in black. (D) Secondary-structure organization of tJ5, consisting of six α-helices: α1 (residues 2-13), α2 (16–20), α3 (25–34), α4 (39–42), α5 (44–47), and α6 (54–63). Helices are represented as red cylinders; random coil regions are shown as black lines. The intramolecular disulfide bonds are shown as connecting lines between the paired cysteine residues. Residue present in the construct are shown in black text. J5 residues absent from the construct are shown in gray, and transmembrane residues are underlined. (E) Topological diagram of tJ5 generated using the PDBsum server. (F) The lowest-energy NMR structure of tJ5 at pH 6.5, as determined by NMR spectroscopy. Ribbon representation of the average NMR structure of tJ5, with α-helices are shown in red and three pairs of disulfide bonds, C10-C19, C21-C46 and C41-C64, highlighted in yellow.

To determine the three-dimensional structure of tJ5, we performed multi-dimensional NMR experiments to obtain information on NOE-derived distance restraints and dihedral-angle restraints for protein structural calculations. During structural refinement, we applied the AMBER force field for energy minimization, incorporating explicit water molecules and ions to approximate the solution environment. These analyses yielded a high-resolution molecular structure with an average root-mean-square deviation (RMSD) of 0.5 Å, as summarized in Table 1. The solution NMR structure of tJ5 consists of six α-helices, α1 (T2-N13), α2 (D16-K20), α3 (D25-D34), α4 (Y39-W42), α5 (E44-K47), and α6 (P54-K63) (Fig. 1C-E). The ectodomain structure is stabilized by three intramolecular disulfide bonds linking α1, α2, α4, α5 and α6: C10-C19, C21-C46, and C41-C64 (Fig. 1F). Notably, helices α1, α3, and α6 form a coplanar triangular ring structure, whereas helices α2, α4 and α5 project outward to form a concave surface. This spatial arrangement suggests that these helices may participate in interactions with other components of the entry-fusion complex (EFC) and thereby contribute to vaccinia virus infectivity.

We next compared the solution NMR structure of tJ5 with the cryo-EM structure of the vaccinia virus EFC. Structural alignment yielded an overall RMSD of 1.176Å across 60 aligned residues (T2-T62), with the largest deviation observed in helices α1 (1.61 Å), α3 (1.37 Å) and α5 (1.33 Å) (Fig. 2A). Consistent with this structural flexibility, J5 engages helices α3 and α4 to interact with A28 protein, whereas helices α1 and α6 contact G9 and A16 within the assembled EFC (43). Specifically, helices α3 and α4 form hydrophobic interactions and hydrogen bonds with adjacent loops of A28 (Fig. 2B, inset I), whereas the α1 and α6 establish hydrophobic contacts and hydrogen bonds with the conserved PXXCW motif of G9 (Fig. 2B, inset II). In addition, the C-terminal region of α6 forms hydrogen bonds with a loop located downstream of the Delta motif of A16 (Fig. 2B, inset III). Together, these α-helical regions form a hydrophobic core while retaining structural flexibility enabling selective protein-protein interactions with A28, G9, and A16 within the vaccinia EFC.

**Figure 2.**
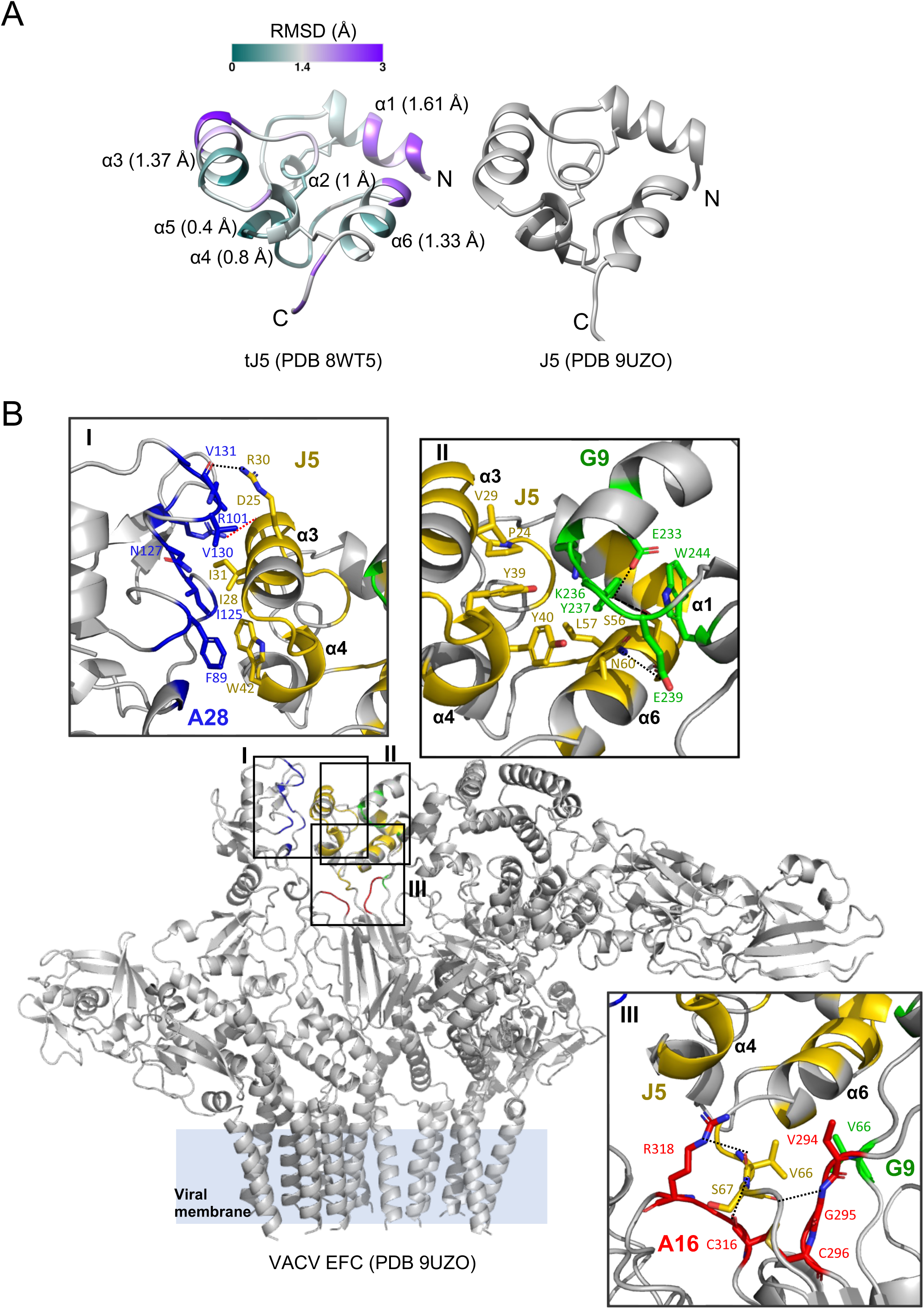
Structural alignment reveals conformational flexibility of α1, α3, α4 and α5 associated with J5 incorporation into the EFC. (A) tJ5 (PDB 8WT5, left) and J5 (PDB 9UZO, right) were structurally aligned in UCSF chimera. RMSD values ranging from 0 to 3 Å, representing increasing structural deviation, are shown as color gradient from teal to gray to purple and mapped onto the tJ5 cartoon model. (B) Cartoon representation of the EFC cryo-EM structure (PDB 9UZO). Residues showing deviations greater than 1.5 Å between tJ5 and cryo-EM J5 are highlighted in yellow. Proteins located within 5 Å of J5 are colored as follows: A28 (blue), G9 (green), and A16 (red). Residues involved in the contact interface are shown as sticks, with hydrogen bonds and salt bridges indicated by dashed lines and colored in black and red, respectively.

### Functional mapping of VACV J5 protein in virus infectivity using recombinant viruses

To determine which residues of J5 are important for vaccinia virus infection, multiple sequence alignment of J5 orthologs from 28 members of the *Poxviridae* family was performed (Fig. 3A). A series of site-directed mutations was generated based on either surface-exposed charged residues (D3, D11, K14) or conserved residues (P^38^YYCWY^43^, S56, N65, N87, Y^103^Q^106^). In addition, all vaccinia J5 orthologues contain eight conserved cysteine residues predicted to form four disulfide bonds (Fig. 3A). We therefore generated a set of five swap mutants, SW(11–18), SW(22–40), SW(42–63), and SW(70–88) each replacing the sequences between cysteine residues, and SW(90–110), which replace the disordered loop before the transmembrane region. (Fig. 3B and Fig. S2A). AlphaFold predicted that each of these SW mutant proteins retained an ectodomain structure (residues 2-68) similar to that of vaccinia WT J5 protein, with RMSD values of 0.886, 1.124, 1.122, 0.933, and 0.957 Å for SW(11–18), SW(22–40), SW(42–63), SW(70–88), and SW(90–110), respectively (Fig. S2B). These results suggest that the chimeric SW constructs likely maintain stable tertiary structures, allowing functional effects to be assessed without global structural disruption. However, SW (70–88) and SW (90–100), which correspond to loop regions in J5 orthologs, showed relatively low prediction confidence (pLDDT <60) and therefore may represent less reliable structural predictions (Fig. S2B).

**Figure 3.**
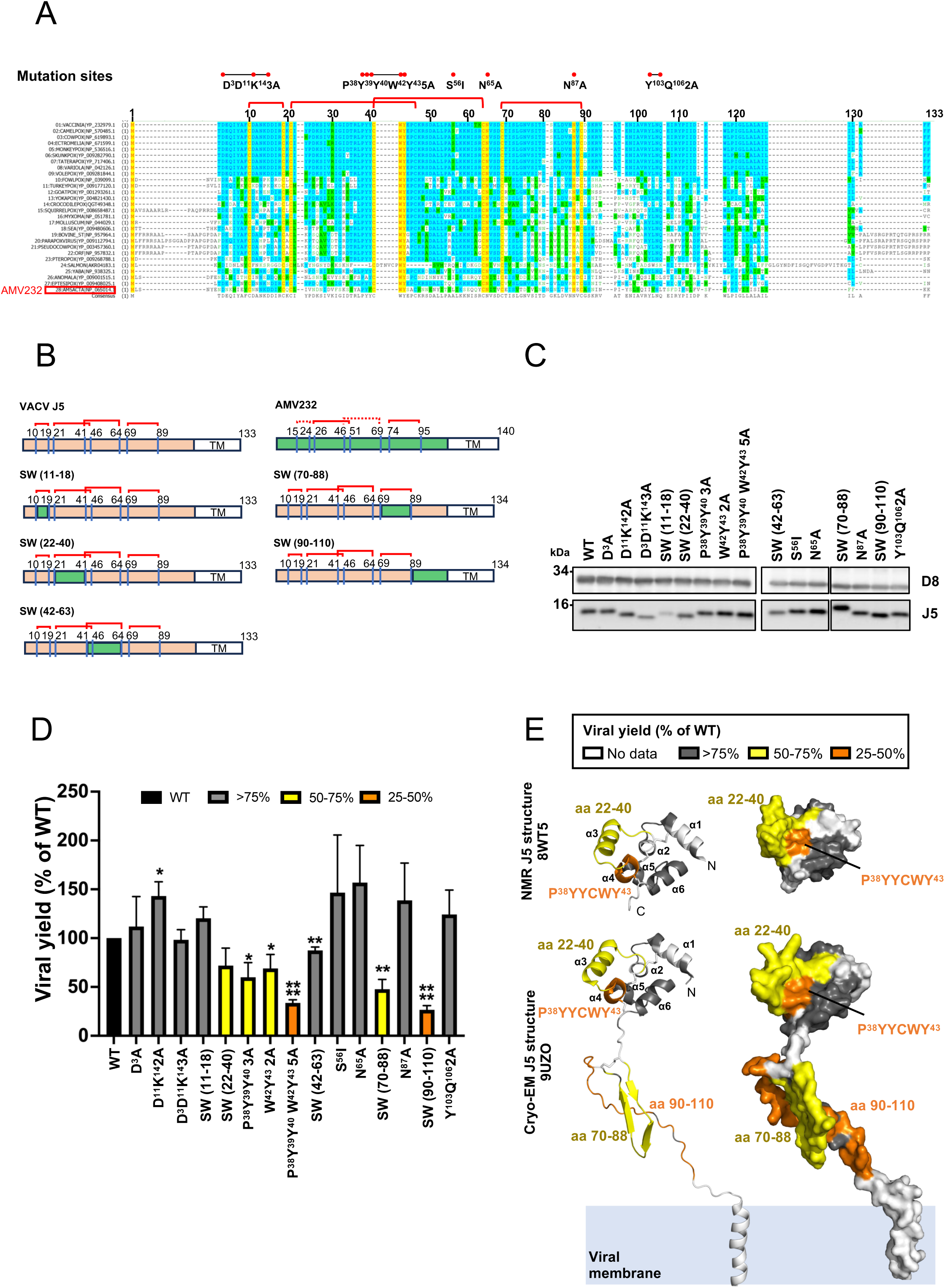
Functional mapping of the vaccinia virus J5 protein. (A) Multiple sequence alignment of J5 orthologues from 28 representative members of the *Poxviridae* family. Identical residues are shown in yellow (100% identity), moderately conserved residues in blue (>50%), and weakly conserved residues in green (>20%). Positions of the targeted mutations are indicated above the alignment. Predicted disulfide bonds are shown as red connecting lines between cysteine residue pairs. (B) Schematic representation of full-length vaccinia virus J5 (residues 1-133, salmon red), AMV232 (residues 1-140, green), and J5 swap mutants (SWs), with chimeric coloring indicating the residue origin. (C) Immunoblot analysis of J5 and the structural protein D8 in infected cell lysates. (D) Virus yields of WT and mutant J5 recombinant viruses in BSC40 cells infected at a multiplicity of infection (MOI) of 5 PFU per cell were harvested at 24 h post-infection and analyzed by plaque assays. Infectivity of each mutant was normalized to that of WT J5. (E) Mapping of virus infectivity loss onto the full-length J5 carton (left panel) and surface (left) models. Regions corresponding to mutants with >75 %, 50-75%, and 25-50% infectivity relative to WT are colored gray, yellow, and orange, respectively. Regions with no experimental data are shown in white. The locations of key functional regions, including residues 22-40, 70-88, and 90-110, are labeled.

Although deletion of *J5L* resulted in a lethal phenotype in vaccinia virus (26, 49), we generated recombinant viruses expressing wild-type (WT) or mutant J5 protein by homologous recombination as described in Materials and Methods. CV-1 cells were infected with WT vaccinia virus and transfected with individual plasmids encoding J5 mutant ORFs together with the *gpt* selection cassette, flanked by the 5’ and 3’ sequences of the *J5L* locus. Recombinant viruses were isolated after three rounds of plaque purification in 1% agar under *gpt* selection, yielding 16 recombinant viruses, including a *gpt*^+^ virus expressing WT J5 protein.

We next evaluated the effect of these mutations on viral infectivity. BSC40 cells were infected at an MOI of 5 PFU/cell, and cell lysates harvested at 24 h post-infection were analyzed by plaque assay and immunoblotting. Expression levels of J5 and the control protein D8 were comparable among most mutants and the WT virus, except for the swap mutant SW(11–18), which showed reduced J5 protein (Fig. 3C). Viral infectivity of WT J5 was normalized to 100% (Fig. 3D). Most J5 mutants retained >75% infectivity (gray bars), including SW(11–18). Four mutants, SW(22–40), P^38^Y^39^Y^40^3A, W^42^Y^43^2A, and SW(70–88), displayed moderate reductions in infectivity (50-75%, yellow bars).

These moderately impaired mutants SW(22–40), P^38^Y^39^Y^40^3A and W^42^Y^43^2A map to helices α3 and α4 of J5 (Fig. 3E), whereas SW(70–88) comprises the Delta motif, a conserved disulfide-bonded closed loop within the A16/G9/J5 protein family that is critical for infectivity and A16/G9 oligomerization (Fig. 3E)(59). Two mutants, P^38^Y^39^Y^40^W^42^Y^43^5A and SW(90–110), exhibited substantial reductions in virus infectivity (33.6% and 26.5%, respectively, orange bars). The P^38^Y^39^Y^40^W^42^Y^43^5A mutant targets the conserved P^38^YYCWY^43^ motif shared among A16, G9, and J5 (P^262^RVCWL^267^ in A16; P^240^RECWD^245^ in G9; P^38^YYCWY^43^ in J5) (59), whereas the SW(90–110) mutant corresponds to a disordered loop located immediately upstream of the transmembrane domain. Together, these infectivity assays indicate that mutations in helices α1, α2, α5 and α6 have relatively minor effects on viral infectivity, whereas mutations in helix α3 and the Delta motif result in moderate reductions in infectivity. In contrast, the P^38^XXCWY^43^ motif and the extended disordered loop spanning residues 90-110 are critical for maintaining EFC function and viral entry efficiency (Fig. 3E).

### Defective J5 mutants with altered P^38^YYCWY^43^ motif and disordered C-terminal loop (90–110) display reduced fusogenic activity

BSC40 cells infected with equivalent amounts of each mutant virus particles were fixed and negatively stained at 24 h post-infection for TEM imaging analyses (Fig. S3). Abundant intracellular mature virion with normal morphology were observed in cells infected with either WT or mutant viruses, consistent with previous findings that repression of EFC components does not affect virion morphogenesis or mature virion formation (18–20, 23–25, 27, 29, 37). We then investigated whether these J5 mutants had lost the ability to trigger membrane fusion, using MV-mediated cell-cell fusion assays as previously described (37, 39). GFP- and RFP-expressing HeLa cells were mixed at a 1:1 ratio and infected with lysates containing comparable amounts of virus particles for 1 h, treated with either neutral or acidic buffer to mimic endosomal acidification, washed and subsequently maintained in growth medium. Cell fusion was monitored every 30 min for up to 3h post-infection. Under neutral pH conditions, little or no fusion was observed for WT or any J5 mutant viruses as expected (Fig. 4A). In contrast, J5 WT induced robust syncytium formation following acid treatment, whereas the P^38^Y^39^Y^40^W^42^Y^43^5A mutant and the SW (90–110) mutant exhibited significantly reduced fusion compared with WT virus (Fig. 4B). Quantifications of virus-induced cell fusion at pH7 and pH5 (Fig. 4C), as well as the relative fusion efficiency of each mutant normalized to WT under low-pH conditions (Fig. 4D), was determined as previously described (61). Together, these data demonstrate that the conserved P^38^YYCWY^43^ motif and the disordered region spanning residues 90-110 are critical determinants required for J5-mediated membrane fusion during vaccinia virus entry.

**Figure 4.**
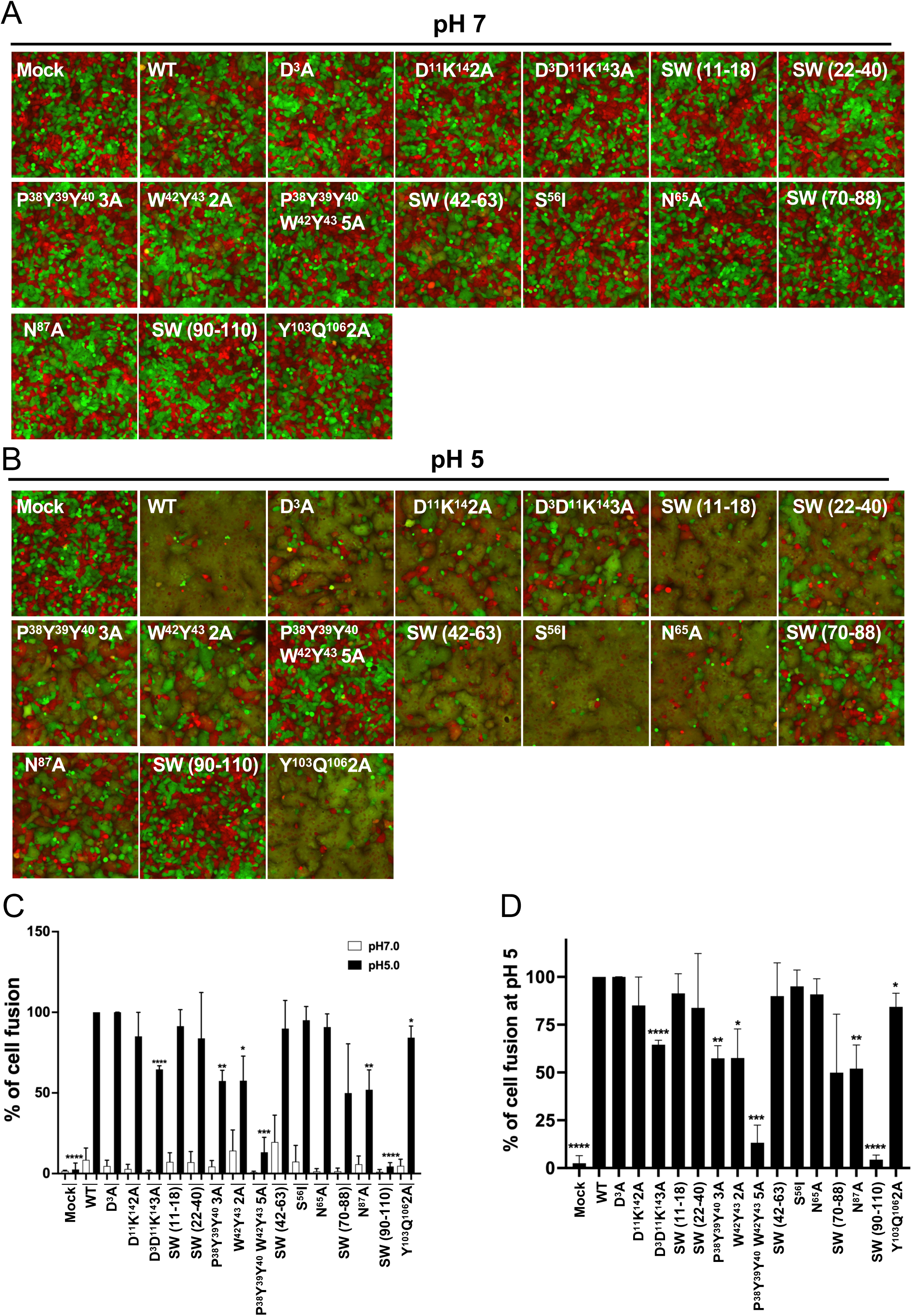
Cell-cell fusion assays of J5 mutants reveal impaired fusion activity at low pH. (A) HeLa cells expressing GFP or RFP were mixed at a 1:1 ratio and incubated with lysates containing WT or mutant J5 recombinant viruses under neutral pH conditions (pH 7). Fluorescence images were photographed at 3 h post-incubation. (B) Parallel fusion assays were performed as described in panel A, except that cells were incubated in acidic buffer (pH 5) for 3 min to mimic endosomal acidification as described in materials and methods. (C) Quantification of MV-triggered cell-cell fusion under neutral or acidic conditions. Images from three independent experiments were analyzed using Fiji software. Fusion was calculated as the percentage of the surface area of double-fluorescent cells relative to that of single-fluorescent cells. White and black bars represent fusion rates at pH 7 and pH 5, respectively. (D) Relative fusion activity of each mutant normalized to WT at low pH. Data represent means ± standard deviations from three independent experiments. Statistical comparison of fusion was performed between J5 WT and each mutant using two-tailed Student’s *t*-test. \**P*<0.05; \*\**P*<0.01; \*\*\**P*<0.001; and \*\*\*\**P*<0.0001.

### The P^38^YYCWY^43^ motif is dispensable for assembly but essential for function, whereas residues 90-110 are required for incorporation of J5 into the EFC

To determine the biochemical basis of the fusion defects, we examined EFC assembly in the two most severely impaired mutants, P^38^Y^39^Y^40^W^42^Y^43^5A and SW (90–110), using co-immunoprecipitation (co-IP). As a control, WT J5 efficiently co-precipitated all EFC components when immunoprecipitated with anti-J5 antibody, confirming that the assay reliably reflects EFC assembly (Fig. 5, left panel). Based on the vaccinia EFC cryo-EM structure, the P^38^YYCWY^43^ motif resides within the helix α4 of J5 and mediates interactions with A28 and G9 through π-π stacking interactions between J5^W42^ and A28^F89^ and between J5^Y40^ and G9^Y237^ (43). Alanine substitution of these aromatic residues was predicted to destabilize these interactions and disrupt complex formation. Unexpectedly, the J5 ^P38Y39Y40W42Y43-5A^ mutant retained interactions with most EFC components (Fig. 5, left panel), and reciprocal co-IP experiments with anti-G9 (Fig. 5, middle panel) and anti-A28 (Fig. 5, right panel) antibodies confirmed that the overall EFC remained intact. Thus, the severe defects in infectivity and membrane fusion observed for the P^38^Y^39^Y^40^W^42^Y^43^5A mutant are not due to disruption of EFC assembly, but instead reflect formation of an assembly-competent yet fusion-inactive EFC.

**Figure 5.**
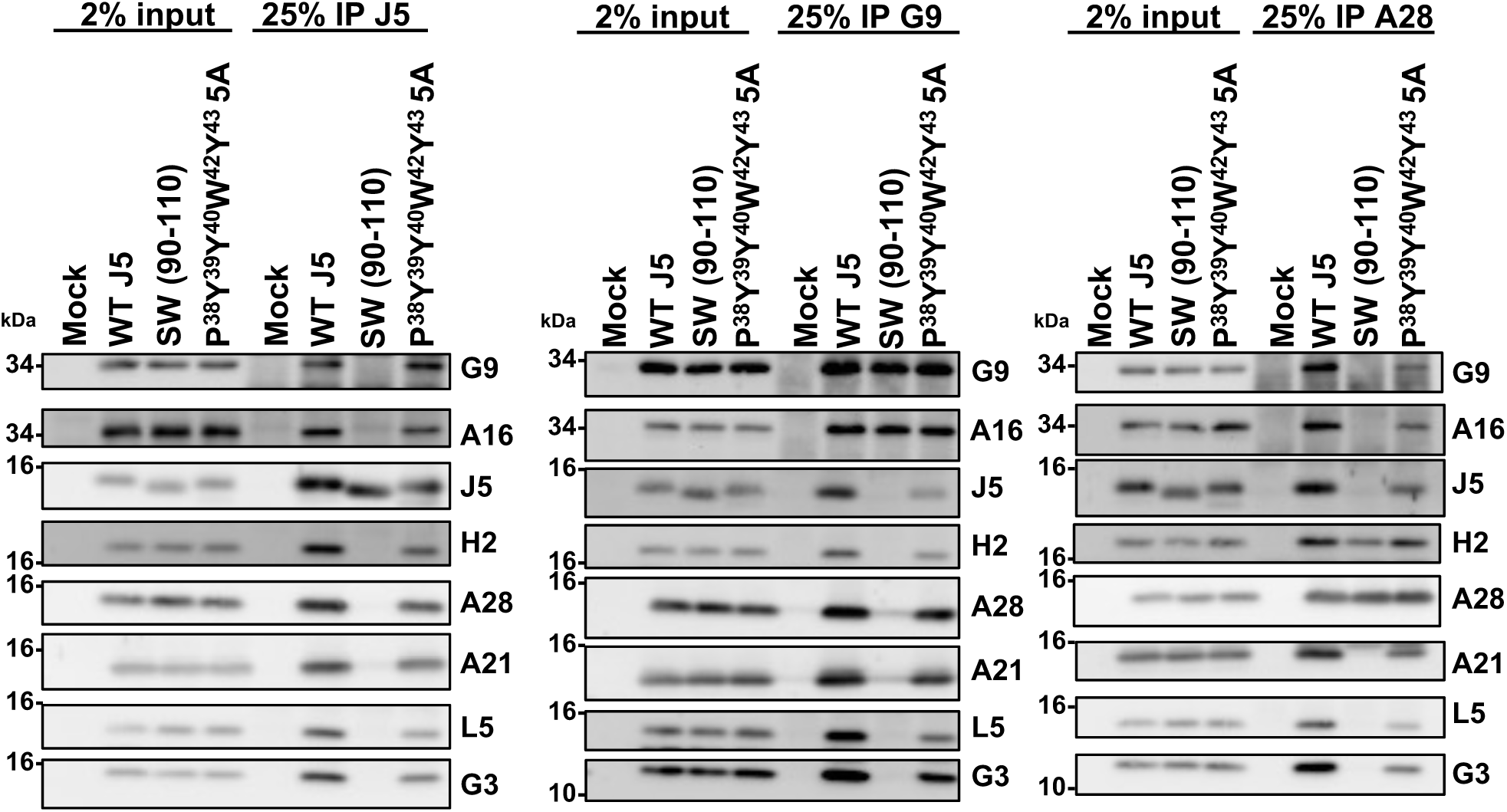
Defective J5 mutants disrupt assembly of the vaccinia virus EFC. BSC40 cells were infected with WT or mutant J5 recombinant viruses. Soluble lysates collected 24 h post-infection were subjected to co-IP using antisera against J5 (left), G9 (middle), or A28 (right), followed by immunoblot analysis with antibodies against EFC components as indicated. The J5 SW(90–110) mutant failed to assemble a complete EFC but retained partial subcomplexes, whereas the J5^P38Y39Y40W42Y43-5A^ mutant exhibited impaired EFC formation.

In contrast, the J5^SW^(^90–110^) failed to co-precipitate A16, G9, A28, H2, A21, L5, and G3 (Fig. 5, left panel). Reciprocal co-IPs using anti-G9 and anti-A28 antibodies showed that the A16/G9 (Fig. 5, middle panel) and A28/H2 subcomplexes (Fig. 5, right panel) remained intact, despite the absence of a full complex assembly. These results indicate that residues 90-110 are specifically required for incorporation of J5 into the central A16/G9/J5 trimer but are not required for the maintenance of individual EFC subcomplexes. Together, these findings reveal two functionally distinct elements within J5: the P^38^YYCWY^43^ motif is not required for EFC assembly but is essential for activation of membrane fusion, whereas the flexible region spanning residues 90-110 is required for stable incorporation of J5 into the EFC.

## Discussion

Vaccinia virus entry relies on the coordinated action of the EFC, a multiprotein machinery that is structurally and mechanistically distinct from classical viral fusion proteins. The cryo-EM structure of the pre-fusion vaccinia EFC was recently solved, revealing that J5 associates with G9 and A16 to form the central trimer, A16/G9/J5(43). In this study, we demonstrate that the conserved P^38^YYCWY^43^ motif located with helix *α*4 and residues 90-110 of J5 are critical for viral infectivity. In addition, mutations in adjacent regions, including residues 22-40 on helix α3 and residues 70-88 corresponding to the Delta motif, also moderately impair EFC function. Mapping these functional sites onto the J5 structure provides a structural framework linking specific regions of J5 to membrane fusion activity of the vaccinia EFC.

In the pre-fusion EFC structure, the P^38^YYCWY^43^ motif of J5 interacts with A28 and G9 through π-π stacking and hydrogen bonding (Fig 6, inset I). Surprisingly, in the fusion-defective P^38^YYCWY^43^5A mutant virus, the EFC remained largely intact, suggesting that P^38^YYCWY^43^ motif contributes to membrane fusion activation rather than EFC assembly. In a recent study, the AlphaFold-predicted model of the EFC suggested substantial rearrangements relative to the cryo-EM pre-fusion structure, particularly in the orientation of the A16/G9 N-terminus and the position of the J5 ectodomain (43). In this predicted model, the J5 ectodomain appears repositioned from the space between A28 and G9 toward the side of A16, which may allow the A16/G9 subcomplex to adopt a more upright conformation. Although the functional relevance of this predicted rearrangement remains unclear, it raises the possibility that the conserved P^38^YYCWY^43^ motif may participate in conformational transitions required for membrane fusion activation. Whether alanine substitution of this motif interferes with such rearrangements remains to be determined.

**Figure 6.**
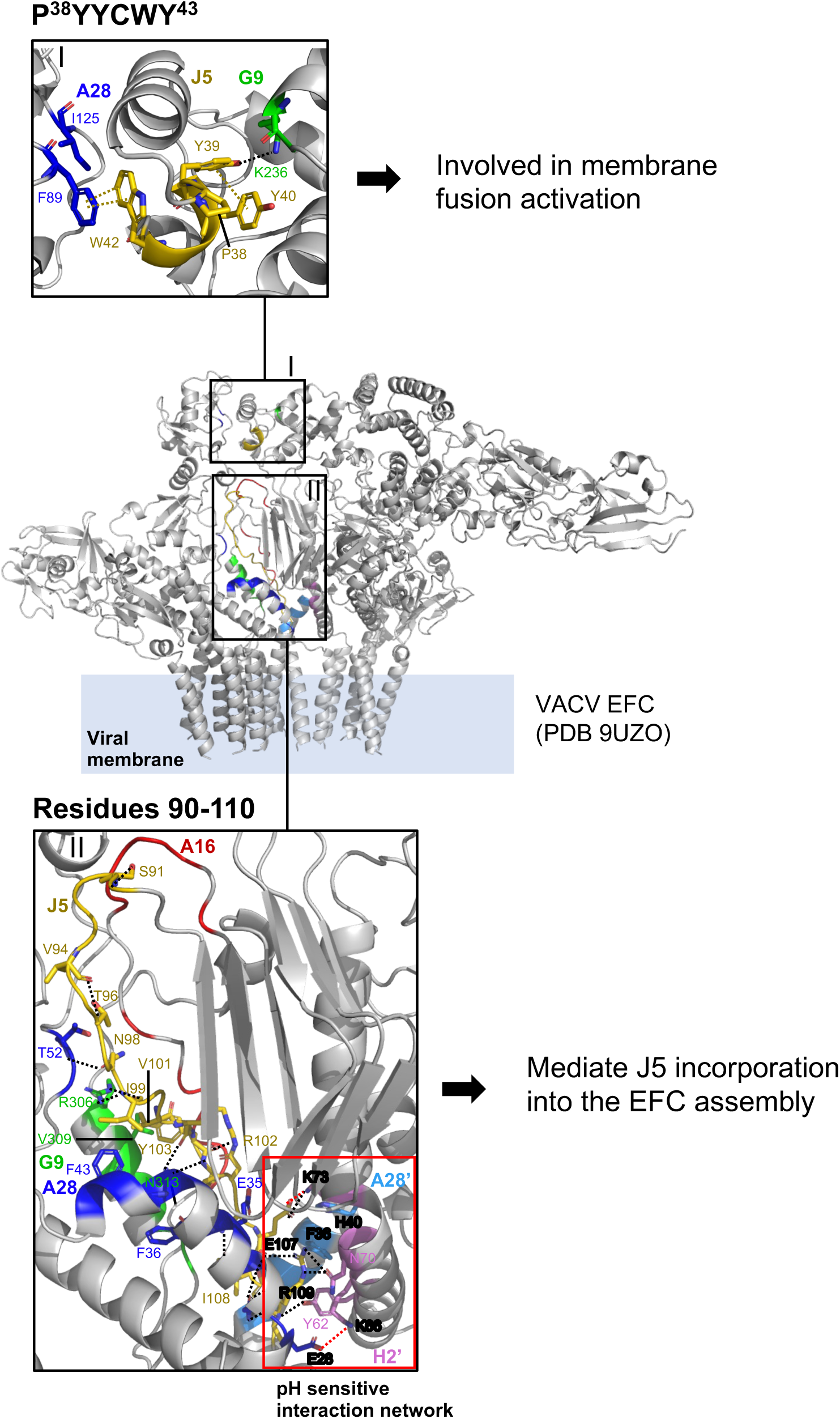
Summary of critical J5 residues and proposed model for EFC assembly. Residue interaction network of the P^38^YYCWY^43^ motif and residues 90-100 of J5 with interacting proteins located within 5 Å in the pre-fusion EFC cryo-EM structure (PDB 9UZO). Interface residues are shown as sticks. Hydrogen bonds, salt bridges, and π- π interactions are indicated by dashed lines and colored black, red, and yellow, respectively.

The second important functional region of J5, residues 90-110, forms extensive contacts with multiple EFC components, including the disordered loop of A16 and the pre-transmembrane helices of G9, A28, A28’ and H2’ (Fig. 6, inset II). These interactions are primarily mediated by hydrogen bonds involving both backbone and sidechain atoms within residues 90-110 of J5 (Fig. 6, inset II, black dashed lines). The extensive contacts with multiple EFC proteins likely explain why the SW (90–110) mutant disrupted EFC assembly, as observed in the co-immunoprecipitation experiments. Notably, we identified a potential pH-sensitive interaction network centered on residues E107 and R109 of J5 (Fig. 6, inset II, red box). At neutral pH, this region is stabilized by two salt bridges, between J5^E107^ and H2’^K73^, and between A28^E28^ and H2’^K66^, thereby reinforcing local contacts among J5, A28, and H2’. In addition, A28’^H40^ lies within 3 Å of H2’^K73^, placing it in close proximity to both H2’^K73^ and J5^R109^. Upon acidification, protonation of histidine residues could render A28’^H40^ positively charged, potentially introducing electrostatic repulsion with H2’^K73^ and J5^R109^. At the same time, protonation of glutamate residues may disrupt salt bridges, weaking local interactions at this interface. Interestingly, a previously reported A28^H40AE44A^ mutant resulted in a loss of viral infectivity and impaired the ability of the EFC to trigger membrane fusion at low pH, which is consistent with this hypothesis (37). Together, these electrostatic changes could destabilize the local interface and potentially facilitate conformational rearrangements of the EFC during membrane fusion. However, whether residues 90-110 of J5 function as a pH-sensitive regulatory element remains to be experimentally tested in the future.

In summary, these findings identified two mechanistically distinct functional elements within J5 that regulate assembly and activation of the vaccinia EFC. The P^38^YYCWY^43^ motif is dispensable for EFC assembly but required for membrane fusion activation, whereas the flexible region spanning residues 90-110 mediates interactions necessary for stable incorporation of J5 into the EFC.

## Acknowledgments

We thank Drs. Chen-Hsin Yu and Hsin-Nan Lin of the IMB Bioinformatics Core and Dr. Shu-Yun Tung of the Genomics Core Facility of the Institute of Molecular Biology, Academia Sinica. NMR spectra were collected at the High-field NMR Center (HFNMRC) supported by Academia Sinica Core Facility and Innovative Instrument Project (AS-CFII-108-112). This work is supported by grants from Academia Sinica, AS-IDR-112-02 (WC) and from National Science and Technology Council, NSTC-113-2320-B-001-016-MY3 (WC) and MOST 111-2113-M-001-014 (DLT).

## Supplementary Figure legends

**Fig. S1. 2D HSQC spectrum of vaccinia virus fusion component tJ5.** 2D ^1^H-^15^N HSQC spectrum of N-isotope-labeled tJ5 (0.2 mM) at pH 6.5. The spectrum was collected at 25 °C in an NMR buffer containing 10 mM phosphate and 137 mM NaCl and 2.7 mM KCl. The assigned residues are indicated using single-letter codes.

**Fig. S2. Design of swap constructs and their predicted 3D models from AlphaFold2.** (A) Sequence alignment of VACV J5 and AMV232. Identical residues are shaded in gray. Predicated disulfide bonds are indicated by red connecting lines. Swap regions between the two proteins are underlined. (B) Cartoon representation of the swap mutant models. J5 Regions containing AMV232 residues are highlighted in green, whereas WT J5 sequences are shown in salmon red (left panel). Predicted local distance difference test (pLDDT) scores are mapped on the chimeric AlphaFold models (right panel). RMSD values relative to the cryo-EM J5 structure are indicated.

**Fig. S3. The normal morphogenesis of J5 WT and J5 mutant viruses.** The BSC40 cells were infected with J5 WT and J5 mutant viruses for 24 h and processed for transmission electron microscopy as described in *Materials and Methods*. Scale bar, 5 mm.

**Fig. S4. Uncropped immunoblot images referred to Figure 3C**. The red boxes indicate the lanes shown in Figure 3C.

